# Identification and characterization of *Gypsophila paniculata* color morphs in Sleeping Bear Dunes National Lakeshore, MI, USA

**DOI:** 10.1101/201350

**Authors:** Marisa L. Yang, Emma Rice, Hailee Leimbach-Maus, Charlyn G. Partridge

**Affiliations:** Environmental Science Policy, and Management, University of California, Berkeley, 260 Mulford Hall, Berkeley, CA 94720; Annis Water Resources Institute, 740 W. Shoreline Dr., Muskegon, MI 49441

## Abstract

**Background:** *Gypsophila paniculata* (baby’s breath) is an invasive species found throughout much of the northwest United States and western Canada. Recently, plants exhibiting a different color morphology were identified within the coastal dunes along eastern Lake Michigan. The common baby’s breath (*G. paniculata*) typically produces stems that are purple in color (purple morph), while the atypical morph has stems that are green-yellow (green-yellow morph). The purpose of this study was to characterize these newly identified morphs and determine if they are genetically distinct species from the common baby’s breath in order to assess whether alternative management strategies should be employed to control these populations.

**Methods:** We sequenced two chloroplast regions, rbcL and matK, and one nuclear region, ITS2, from the purple morphs and green-yellow morphs collected from Sleeping Bear Dunes National Lakeshore, MI, USA (SBDNL). Sequences were aligned to the reference sequences from other *Gypsophila* species obtained from the Barcode of Life (BOLD) and GenBank databases. We also collected seeds from wild purple morph and wild green-yellow morph plants in SBDNL. We grew the seeds in a common garden setting and characterized the proportion of green-yellow individuals produced from the two color morphs after five-months of growth.

**Results:** Phylogenetic analyses based upon rbcL, matK, and ITS2 regions suggest that the two color morphs are not distinct species and they both belong to *G. paniculata*. Seeds collected from wild green-yellow morphs produced a significantly higher proportion of green-yellow individuals compared to the number produced by seeds collected from wild purple morphs. However, seeds collected from both color morphs produced more purple morphs than green-yellow morphs.

**Discussion:** Based upon these results, we propose that the two color morphs are variants of *G. paniculata*. Given the significant difference in the number of green-yellow morphs produced from the seeds of each morph type, we also suggest that this color difference has some genetic basis. We propose that current management continue to treat the two color morphs in a similar manner in terms of removal to prevent the further spread of this species.

## Introduction

The Great Lakes sand dunes comprise the most extensive freshwater dune complex in the world, stretching over 1,000 km^2^ in Michigan alone. Within northwest Michigan, the sand dunes ecosystem is vital both environmentally and economically. It is home to a number of threatened and endangered species, including piping plover (*Charadrius melodus*) and Pitcher’s thistle (*Cirsium pitcheri).* Colonization of invasive species in this region has the potential to significantly alter the biological composition of these native communities (Leege and Murphy, 2001; Emery et al. 2013). One invasive species of significant concern is the perennial baby’s breath (*Gypsophila paniculata*). In 2015, baby’s breath was listed by the Michigan Department of Natural Resources (DNR) as a “priority” invasive species for detection and control in Michigan’s northern lower peninsula (DNR, 2015). Since its colonization in the region it has spread along a 260 km stretch of the Michigan shoreline. Baby’s breath produces a large taproot system that can extend down to 4 meters in depth, which likely helps it outcompete native vegetation for limited resources (Darwent & Coupland, 1966; Karamanski, 2000). In addition, while many of the vulnerable and endangered plant species in these areas are seed limited, (e.g., Pitcher’s thistle produces approximately 50-300 seeds per plant total or ‘per lifetime’ (Bevill et al., 1999)), baby’s breath can produce up to 14,000 seeds per plant annually (Stevens, 1957), effectively outcompeting native species in terms of overall yield. This has led to baby’s breath composing approximately 50-80% of the ground cover in some areas (Karamanski, 2000; Emery et al., 2013).

One concern with current management efforts is that anecdotal evidence suggests there may be a new baby’s breath variant within the Michigan dune system. In 2011 and 2012 The Nature Conservancy (TNC) removal crews reported baby’s breath plants with different character traits than what is commonly observed (TNC, 2014). The atypical morph has stems and leaves that are lighter in color and more yellow than the common *G. paniculata* purple morph (Figure 1a-c). The purple morph has a thick taproot (4-7 cm in diameter) just below the caudex that remains unbranched for approximately 60 – 100 cm (Darwent & Coupland, 1966). Severing just below the intersection of the caudex and the taproot is where manual removal efforts target to limit regrowth. However, TNC removal crews suggested that the atypical green-yellow morph’s root system seemed to be more diffuse, making it harder to identify a primary taproot and thus, harder to sever without the potential for regrowth (TNC, 2014). Currently, these green-yellow morphs are treated with herbicide application (glyphosate) when observed; however, if this is a newly invaded baby’s breath species and it continues to spread into areas where threatened or endangered species are present, removal methods will be a primary concern and alternative management strategies may need to be considered for these populations.

**Figure 1:**
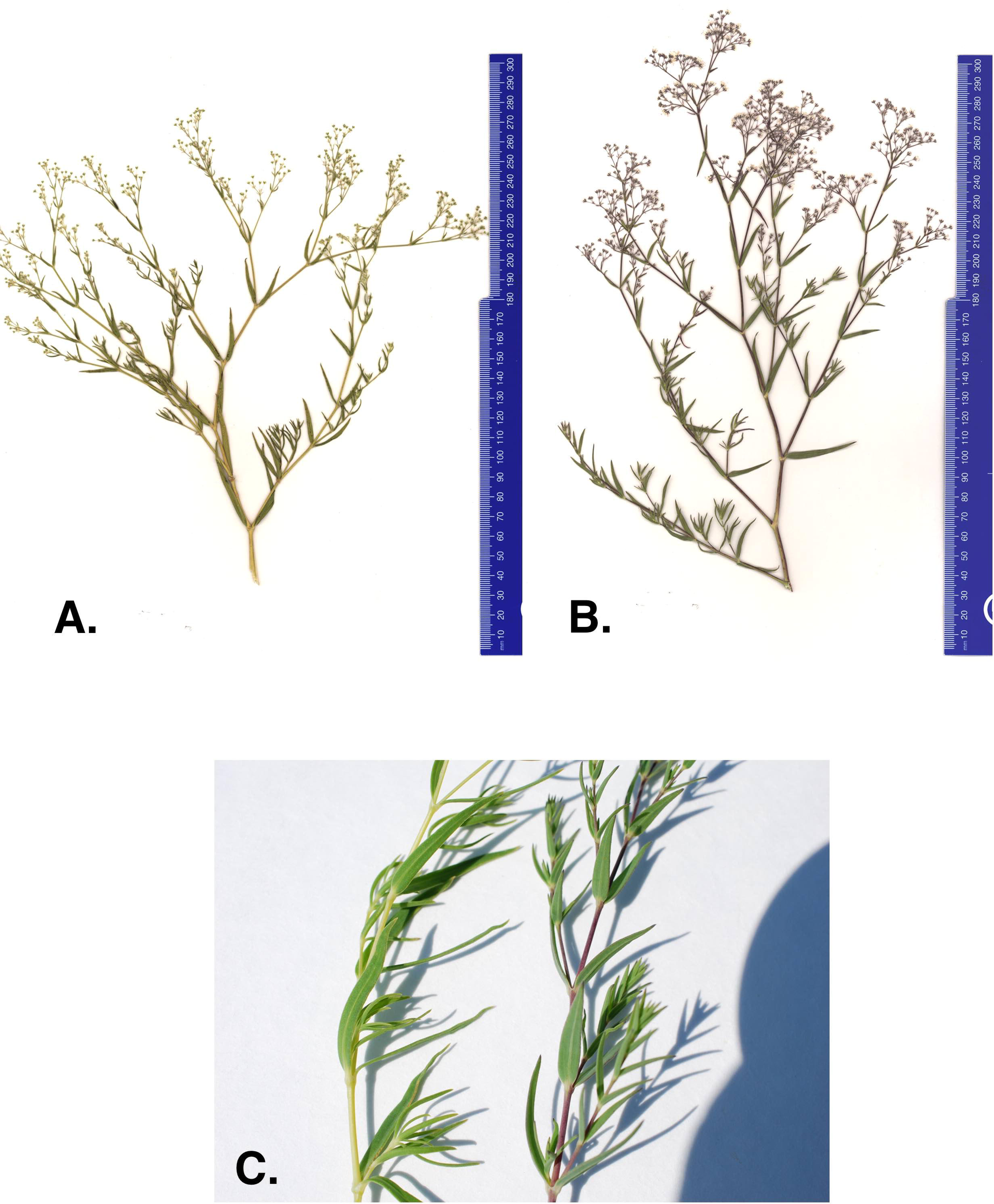
(A) Green-yellow morph, (B) common purple morph and (C) stem of the green-yellow and purple baby’s breath morph found in Sleeping Bear Dunes National Lakeshore.

One of the first steps toward adapting current management strategies for this invasive is to identify whether the green-yellow morph is a genetically distinct species from the purple morph. While *G. paniculata* is the dominant invasive baby’s breath species in northwest Michigan, a number of other species have been introduced to North America and, specifically, the Great Lakes region (Pringle, 1976, Voss & Reznicek, 2012). For example, *G. elegans, G. scorzonerifolia, G. muralis,* and *G. acutifolia* have been collected within Michigan (Pringle, 1976; Reznicek et al., 2011; Voss & Reznicek, 2012), and *G. perfoliata* is reported to have become naturalized in the United States (Pringle, 1976). *G. muralis* is an annual and has a very distinct morphology compared to the other *Gypsophila* species identified around the Great Lakes. It typically only reaches 5-20 cm in height, has linear leaves, and commonly produces white to pink flowers (Borkaudah, 1962). *G. elegans*, also an annual, is commonly sold in this region in commercial wildflower packets. It typically has a smaller taproot compared to *G. paniculata,* and its coloration can be similar to that observed for the green-yellow morph*. G. scorzonerifolia* and *G. acutifolia* are perennials and specimens of these species have been collected in counties within the Great Lakes dune system that also contain *G. paniculata* infestations (Voss, 1957; Pringle, 1976). Both *G. scorzonerifolia* and *G. acutifolia* have a deep taproot and are similar in height to *G. paniculata,* and thus, can superficially resemble *G. paniculata* (Voss, 1957; Pringle, 1976). However, both can be distinguished from *G. paniculata* in that their leaves tend to be longer and wider, and the pedicels and calyces are glandular as opposed to glabrous in *G. paniculata* (Voss, 1957; Pringle, 1976). Given the potential for these other species to invade the fragile habitat of the Michigan dune system, the goal of this work was to characterize the genetic relationship between the newly recognized green-yellow morph and the common purple morph to determine if they are the same species.

## Methods and materials

### DNA Extraction, Amplification, and Sequencing

We collected leaf tissue from 1 green-yellow morph and 16 purple morphs in 2016 and an additional 15 green-yellow morphs in 2017 from Sleeping Bear Dunes National Lakeshore (SBDNL), Empire, MI, USA (specifically: 44.884941 N, 86.062111 W and 44.875302 N, 86.056821 W). Plant tissue collections were approved by the National Parks Service (permit ID SLBE-2015-SCI-0013). Leaf tissue was dried in silica gel until DNA extractions could take place. DNA was extracted using a Qiagen DNeasy Plant Mini Kit (Qiagen, Hilden, Germany). After extraction, the DNA samples were placed through Zymo OneStep PCR inhibitor removal columns (Zymo, Irvine, CA) to remove any secondary metabolites that might inhibit PCR amplification. The DNA for each sample was then quantified using a NanoDrop 2000 (ThermoFisher, Waltham, MA).

The DNA of green-yellow morphs and purple morphs was amplified at three genetic regions: large subunit of the ribulose-bisphosphate carboxylase gene (rbcL), maturase K (matK), and internal transcribed spacer 2 (ITS2). A combination of ITS region and matK have been used to differentiate between other *Gypsophila* species (specifically, *G. elegans* and *G. repens*) in previous studies (Fior et al., 2006). The rbcL region was amplified using rbcL 1F and rbcL 724R primers (Fay et al., 1997), matK was amplified using matK 390F and matK 1440R primers (Fior et al., 2006), and the ITS2 region was amplified using ITS2 2SF and ITS2 S3R primers (Chen et al., 2010). PCR reactions for all loci consisted of 1X Taq Buffer, 2.0 mM MgCl_2_, 0.3 μM dNTP, 0.08 mg/mL BSA, 0.4 μM forward primer, 0.4 μM reverse primer, and 0.5 units of Taq polymerase in a 20 μL reaction volume. The thermal cycle protocols consisted of the following: for rbcL, an initial denaturing step of 95°C for 2 minutes, followed by 35 cycles of 94°C for 1 minute, 55°C for 30 seconds, and 72°C for 1 minute. A final elongation step was performed at 72°C for 7 minutes. For matK, the thermal profile consisted of 26 cycles of 94°C for 1 minute, 48°C for 30 seconds, and 72°C for 1 minute, followed by a final elongation step at 72°C for 7 minutes. For ITS2, an initial denaturing step of 95°C for 2 minutes was applied, followed by 35 cycles of 95°C for 30 seconds, 50°C for 30 seconds, 72°C for 1.5 minutes, and a final elongation step of 72°C for 8 minutes. Successful amplification was checked by running the PCR product on a 2% agarose gel stained with ethidium bromide. PCR reactions were then cleaned using ExoSAP-IT PCR Product Cleanup Reagent (ThermoFisher, Waltham, MA). Sequencing reactions were performed with the forward and reverse primers for each of the three regions. Sequencing reactions were cleaned using a Sephadex column (GE Healthcare Life Science, Marlborough, MA) and sequenced on an ABI Genetic BioAnalyzer 3130xl (Applied Biosystems, Foster City, CA). Out of the 16 green-yellow morphs a total of 13 were successfully sequenced for rbcL, 13 were successfully sequenced for matK, and 14 were successfully sequenced for ITS2. For the purple morphs a total of 15, 12, and 15 individuals were successfully sequenced for rbcL, matK and ITS2, respectively.

Reference sequences for rbcL, matK, and ITS2 of other *Gypsophila spp*. were downloaded from either the Barcode of Life Database (BOLD) (http://www.barcodeoflife.org) or GenBank (https://www.ncbi.nlm.nih.gov/genbank/). We primarily focused on *Gypsophila* species with reported occurrences within the United States, but also incorporated other species if their information was available on BOLD. Sequences of the three regions were not always available for the same species, thus for rbcL these included *G. paniculata, G. elegans, G. fastigiata, G. scorzonerifolia, G. perfoliata*, and *G. murali*s. For matK the species included *G. paniculata, G. elegans, G. fastigiata, G. scorzonerifolia, G. perfoliata, G. muralis, G. altissima*, and *G. repens*. For the IST2 region, the species included *G. paniculata, G. elegans, G. scorzonerifolia, G. perfoliata, G. repens,* and *G. acutifolia.* Because sequences for all three regions were not available for all species, our merged phylogeny only contained *G. paniculata, G. elegans, G. scorzonerifolia,* and *G. perfoliata* reference sequences. The accession numbers and sequences for all reference species are provided in Supplemental Table 1. All FASTA files corresponding to these data will be deposited in the Dryad database and sequences have been submitted to GenBank (ITS2: MG385003-385031, matK: MG603322-603346, rbcL: MG547346-547373).

### Alignment and Phylogenetic Analysis

All successful sequences from our field samples, as well as sequences for other *Gyposphila* species reference sequences obtained from BOLD or GenBank, were imported into the program MEGA7 (version 7.0.14) (Kumar et al., 2016) and sequences for each of the three regions were aligned both individually and with all sequences combined using Muscle (Edgar, 2004). The total number of base pairs aligned and analyzed for each region included: 427 base pair (bp) for rbcL, 702 bp for matK, 201 bp for ITS2, and 1330 bp for the three regions combined. All alignment parameters were kept at their default settings. Once aligned, we used MEGA7 to identify the most appropriate substitution model (rbcL: Jukes-Cantor, matK: Tamura 3-parameter, ITS2: Jukes-Cantor, all genes combined: Tamura 3-parameter with gamma distribution). We then created phylogenetic trees using a maximum-likelihood (ML) approach with 500-replicated bootstrap analyses, as well as using neighbor joining, and parsimony models. We also constructed a TCS haplotype network (Clement et al., 2002) based upon the combined sequences using the statistical parsimony approach (Templeton et al., 1992) in the program PopART (v 1.7, http://popart.tago.ac.nz).

### Color Morph Germination

On May 8, 2018 we planted a total of 207 seeds collected from mature purple morphs and 255 seeds collected from mature green-yellow morphs from SBDNL. For the purple morph seeds, these were collected from a total of 14 plants that were sampled in 2016 (average 15 seeds per plant) and 7 plants that were sampled in 2017 (average 1.7 seeds per plant). For the green-yellow morphs, these seeds were collected from a total of 17 plants (15 seeds per plant) in 2017. Plants were grown in the GVSU – Allendale greenhouse from May until August, 2018. The greenhouse was on a 17 hour light/ 7 hour dark cycle. The average day temperature was 21°C and night temperature was 15 °C. In August the plants were transported to the greenhouse at AWRI-GVSU where they were allowed to grow until October 15, 2018. The greenhouse at AWRI-GVSU has no external lighting source or temperature controls, and thus more closely resembled seasonal day/night and temperature cycles. Plants were sampled after a decrease in temperature occurred (from a high of 23 °C on October 10, 2018 to a high of 11 °C on October 15, 2018). Previous greenhouse observations have found that the differences between the purple and green-yellow morphs can be best detected after a sudden drop in temperature (personal observation, CGP). On October 15, we characterized the color of all individuals that successfully germinated and survived over the 5-month period. We used a chi-square analysis in the R statistical package (v3.5.1) to determine if germination success differed between seeds from the two color morphs and whether the proportion of seeds that developed into green-yellow morphs significantly differed between seeds collected from mature purple and mature green-yellow plants.

## Results

Our results indicate that the green-yellow morph identified in SBDNL is not a genetically distinct species from the common purple found throughout SBDNL. The rbcL, matK, ITS2, and combined dataset showed similar patterns with both the green-yellow morphs and the purple morphs clustering together. The phylogenies constructed from rbcL and matK independently show that the two color morphs cluster separately from *G. fastigata, G. elegan, G. muralis*, and *G. repens*. For the rbcL locus, the relationship of the color morphs to *G. paniculata* and *G. scorzonerifolia* was not resolved. Additionally, when we only examined the matK gene, the color morphs clustered separately but within a clade that also included *G. altissima, G. scorzonerifolia*, and *G. paniculata*. The ITS2 region was able to provide more resolution between *G. paniculata, G. scorzonerifolia, G. acutifolia*, and the color morphs, with the color morphs clustering with *G. paniculata* (Figure 2a-d; Supplemental Figure 1-8) and separately from the *G. scorzonerifolia* and *G. acutifolia* clade. The same pattern was observed when all regions were analyzed together (Figure 2d), with the exception that this phylogeny did not include *G. acutifolia*. In addition, the TCS haplotype network shows that the purple and green-yellow morphs have shared haplotypes. These two haplotypes are only one mutation away from one another and the *G. paniculata* reference, while the next closest species, *G. scorzonerifolia* is 15 mutations away (Figure 3). This further suggests that both color morphs are *G. paniculata*.

**Figure 2:**
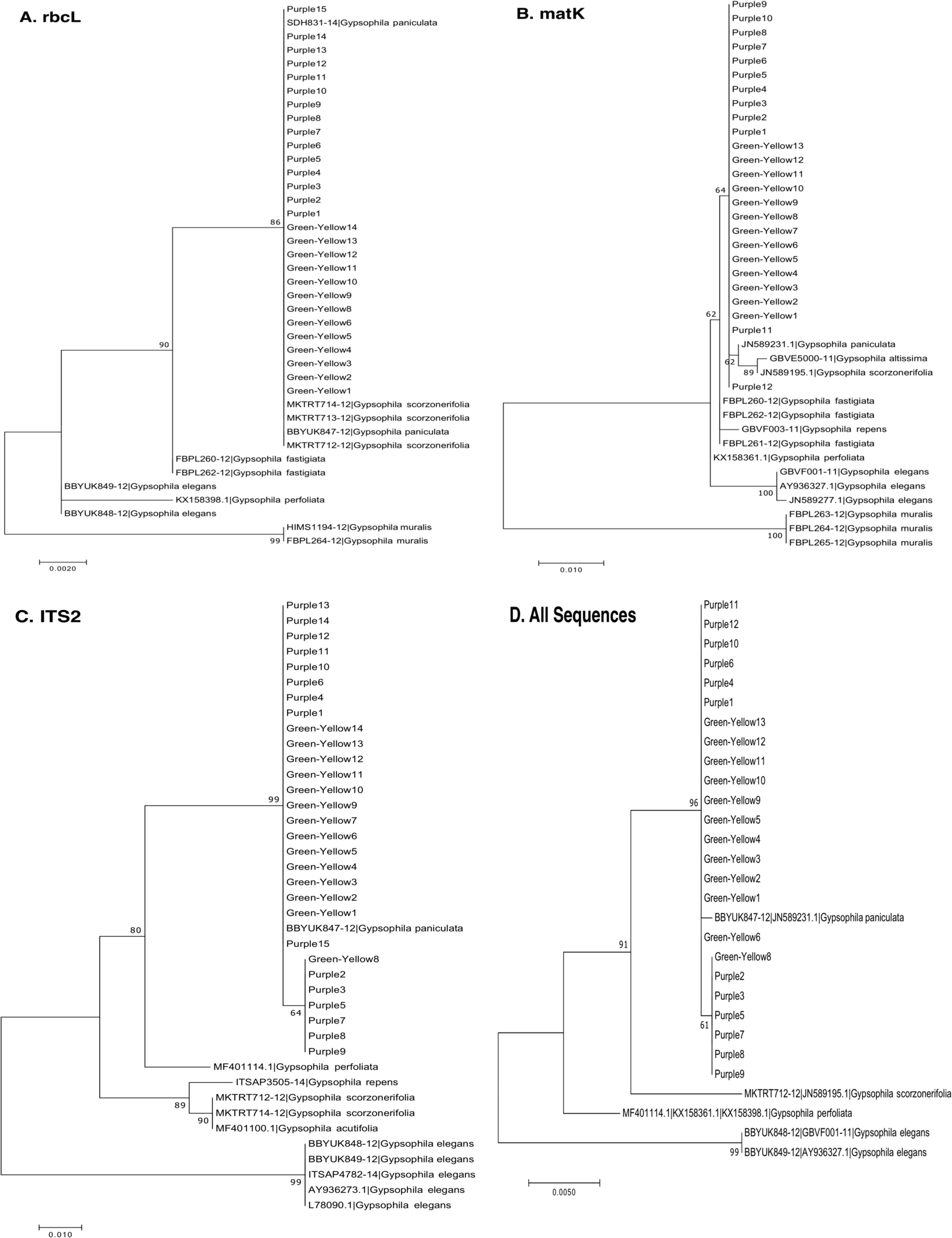
Phylogenetic analysis of the purple and green-yellow baby’s breath color morphs in relationship to other *Gypsophila* species. All evolutionary histories were inferred using maximum likelihood methods. (A) Phylogeny based on the rbcL region, (B) phylogeny based on the matK region, (C) phylogeny based on the ITS2 region, (D) Phylogeny based on rbcL, matK, and ITS2 combined. For rbcL and ITS2 we used a Jukes Cantor (JC) model of molecular evolution (Jukes and Cantor, 1969). For matK we used a Tamura 3-parameter (T92) model of molecular evolution with uniform distribution, and for the combined data set we used the T92 model of molecular evolution with a gamma distribution (Tamura 1992).

**Figure 3:**
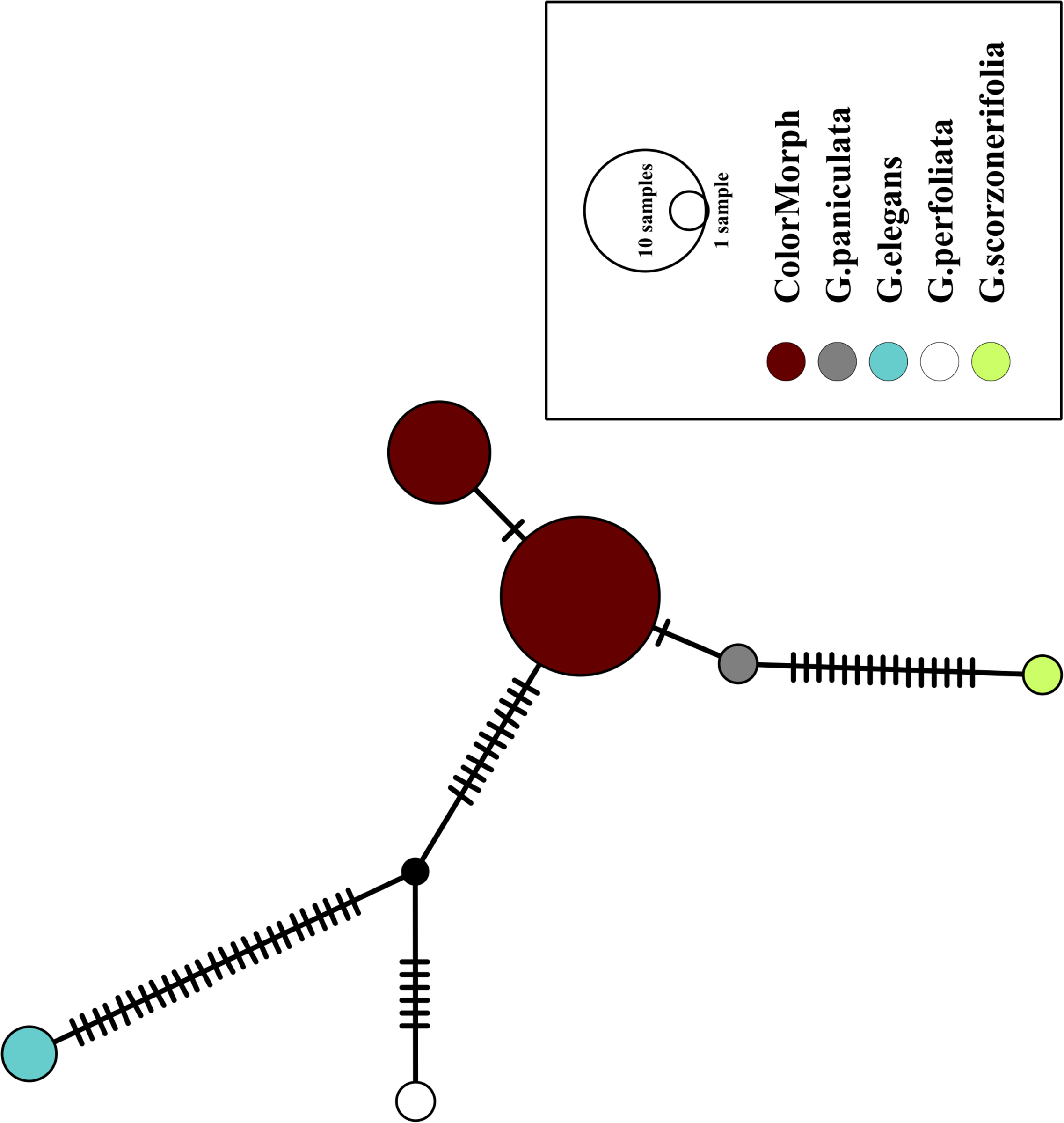
A TCS haplotype network based on rbcL, matK and ITS2 combined for the purple and green-yellow baby’s breath color morphs and the *G. paniculata, G. elegans, G. perfoliata*, and *G. scorzonerifolia* reference sequences. The size of the ovals correspond to the haplotype frequency. The hash marks represent the number of mutations between each haplotype.

Of the regions analyzed for the green-yellow morphs, purple morphs, and reference sequences, rbcL was the most conserved sequence with an overall mean genetic distance (*d)* = 0.004, followed by matK (*d* = 0.015) and ITS2 (*d* = 0.038). For the ITS2 region, there were six purple morphs and one green-yellow morph that clustered together inside the *G. paniculata* branch (Figure 2c & 2d). Further examination of the electropherograms for these individuals show that they are likely heterozygous at position 138 of our aligned sequence and amplification bias of the ‘A’ single nucleotide polymorphism (SNP) over the allele containing the ‘G’ SNP is driving this pattern.

### Color Morph Germination

Out of the 207 seeds that were collected from mature purple morphs and planted in the greenhouse, 82 successfully germinated and survived over the 5 month period (39.6%). Out of these 82 plants, only one green-yellow morph was produced (1.2%), while the remaining seeds all produced purple morphs. Out of the 255 seeds collected from mature green-yellow morphs and planted in the greenhouse, 105 successfully germinated and survived over the 5 month period (41.2%). This was not significantly different than the proportion of plants that successfully germinated from the purple morph seeds (χ^2^ = 0.06, df = 1, p = 0.81). Of the 105 successfully germinated seeds from the green-yellow morph plants, 12 developed into green-yellow morph plants (11.4%), 91 developed into purple morphs plants (86.7%), and two plants could not be determined (they appeared to be green-yellow morphs but displayed some dark spots on the stem). The proportion of seeds that produced green-yellow individuals significantly differed between seeds collected from mature green-yellow morphs and seeds collected from mature purple morphs (χ^2^ = 5.9, df = 1, p = 0.015).

## Discussion

Overall, our data suggest that the green-yellow morph is not a genetically distinct species from the purple morph, and that both morphs are *G. paniculata*. For all molecular markers used, the green-yellow and the purple color morphs grouped together. RbcL, matK, and ITS2 are common ‘barcode’ genes used to delineate plant species (Newmaster et al., 2006; Group et al., 2009; Chen et al., 2010; Yao et al., 2010; Stoeckle et al., 2011) and when used in combination they provided adequate resolution to separate out the *Gypsophila* species included in this study. In our data set, rbcL and matK worked well to separate our color morphs from *G. elegans, G. muralis* and both of these species have been reported to occur in the Great Lakes region (Reznicek et al., 2011; Voss & Reznicek 2012). While the morphology of *G. muralis* is very distinct from the color morphs in SBDNL, it was initially thought that *G. elegans* shared some similar traits to that originally described by the TNC removal crews and was a potential candidate species for the green-yellow color morph. Based upon these results, this is clearly not the case.

The phylogeny based on ITS2 region and the combined sequences provided the best resolution for assigning the relationship of our *Gypsophila* species. Like the rbcL and matK phylogenies, all the purple and green yellow morphs grouped together. For this region, the color morphs also grouped within the same clade as the *G. paniculata* reference sequence. While *G. scorzonerifolia* and *G. acutifolia* have also been recorded in the Great Lakes regions (Pringle, 1976), and have a similar general phenotype as *G. paniculata*, these species were clearly within a distinct clade that was separate from the two color morphs. Similarly, while *G. perfoliata* has been reported to be naturalized in North America (Pringle, 1976), it grouped outside of the *G. paniculata* and color morph cluster.

Our greenhouse germination study showed that seeds collected from mature green-yellow morphs produced a significantly higher proportion of green-yellow individuals than seeds collected from mature purple morphs. Of the seeds collected from mature green-yellow morphs 11% resulted in green-yellow morphs, while only 1% of seeds from mature purple morphs resulted in green-yellow morphs. However, seeds from both color morphs primarily produced purple morphs. The mechanism driving the color difference between the purple and green-morphs is currently unknown. Within SBDNL, the purple morph is the most common form, with green-yellow individuals found interspersed in a couple of locations throughout the dunes (personal observation, CGP/HLM/ER). The largest observed group of green-yellow morphs consists of a few hundred plants clumped within approximately an acre-sized area and interspersed throughout large groups of purple morphs. Based upon the dispersal patterns of the two morphs throughout the dunes, and our germination results, the color difference observed does not appear to be solely environmentally driven, and likely has a genetic component. Potential candidate genes that could be influencing these color differences include those involved in the anthocyanin pathway, which influences red – purple coloration in a number of plants (Asen et al., 1972; de Pascual-Teresa et al., 2002; Abdel-Aal et al., 2006). In addition, anthocyanin can rapidly accumulate in the shoots of plants following cold exposure (Leng et al., 2000), and we have observed an increase in the amount of purple coloration in the purple morphs as temperatures decrease (personal observation, CGP). Further work will begin to elucidate the specific mechanism influencing this color difference in invasive *G. paniculata* populations, as well as to explore whether this color variation drives functional differences between the morphs.

Taken together, these data show that the purple and green-yellow morph within SBDNL are the same species, and that species is *G. paniculata*. One concern with the green-yellow morph initially noted by TNC removal crews was that the taproot tended to be more diffuse than the purple morph, potentially making manual removal of these plants less effective. However, we have not noted differences in the taproot structure between these two morphs when grown under controlled conditions (Figure 4). Additionally, our lab’s personal observations (CGP, HLM) in the field have not found any indication that large differences in root structure occur between mature plants of the two color morphs. Therefore, current management approaches for these populations should be maintained to control the further spread of *G. paniculata* throughout the Michigan coastal dune system.

**Figure 4:**
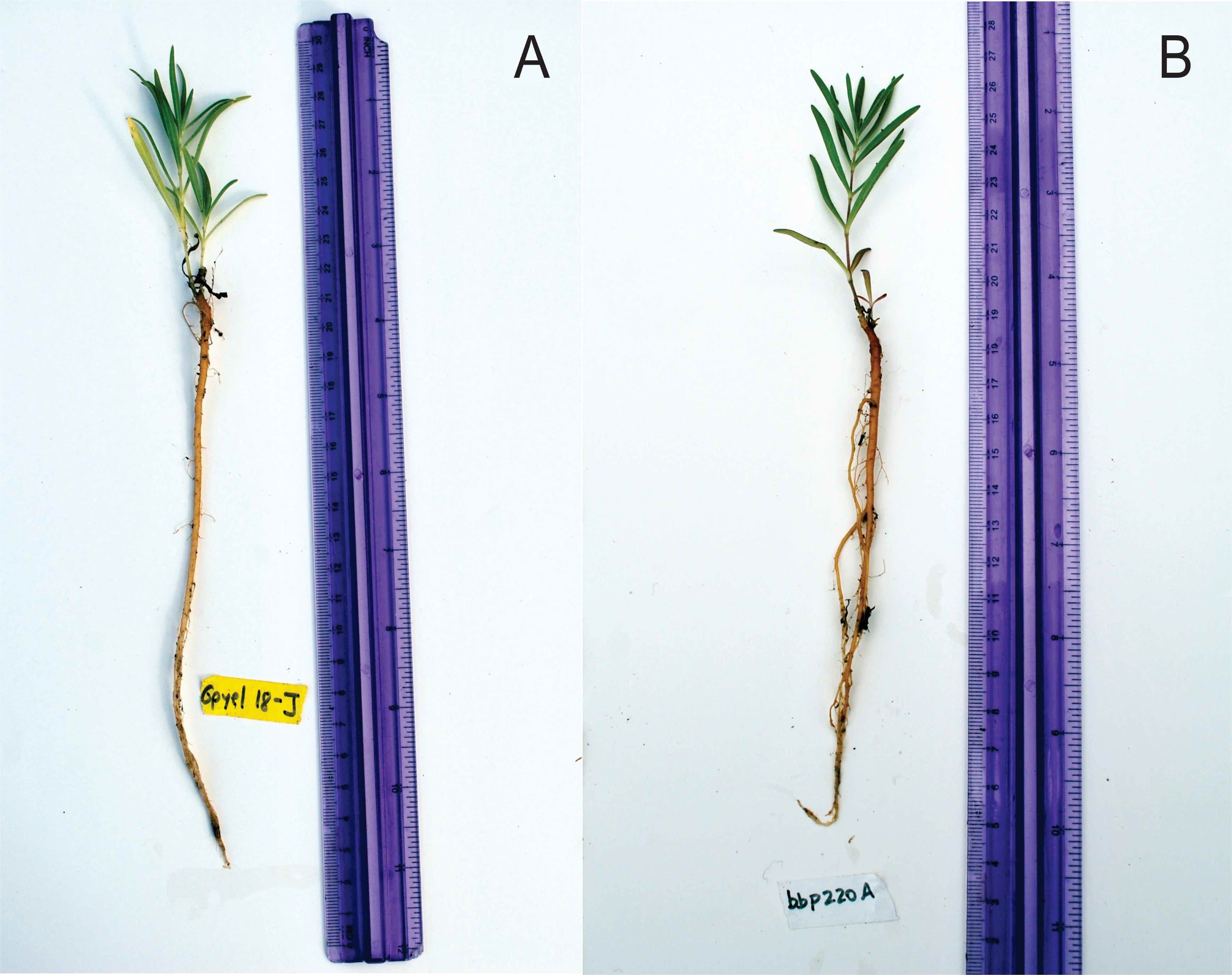
(A) Green-yellow morph, and (B) common purple morph after 5-months in the GVSU greenhouse. Note the similarity in taproot structure between the two plants.

## Conclusions

Our data show that both the purple and green-yellow color morph of baby’s breath in Sleeping Bear Dunes National Lakeshore are *G. paniculata* and the observed color differences likely have some genetic basis. Based on this current information, we recommend that these color morphs continue to be managed in a similar manner and that distinct management strategies do not need to be established at this time.

## Supporting information

Raw sequence files, alignment files

Gypsophila reference sequences

Supplemental phylogenies

## Acknowledgements

We would like to thank Shaun Howard from The Nature Conservancy for his help in identifying the green-yellow morph and Benjamin Giffin for his help with sequencing analysis. We would also like to thank Kurt Thompson and Doug Haywick for assisting with the figure construction, and Alexis Hoskins and Doug Haywick for helping with planting. Funding for this project was provided through an EPA-Great Lake Restoration Initiative Grant and an NSF-Research Experience for Undergraduates Grant.

