## Supplemental phylogenies for "Identification and characterization of *Gypsophila paniculata* color morphs in Sleeping Bear Dunes National Lakeshore, MI, USA"

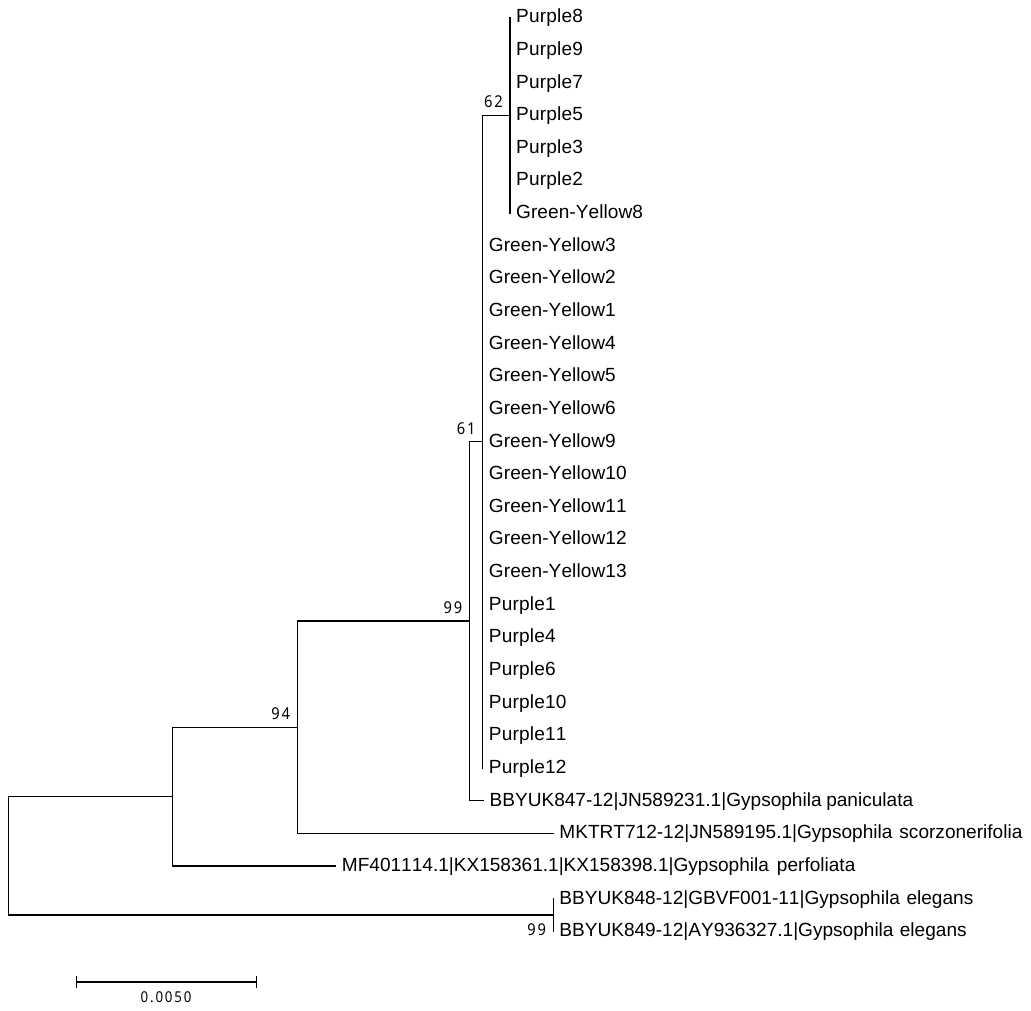

**Supplemental Figure 1.** **Neighbor-Joining analysis of baby’s breath color morphs in relation to other *Gypsophila* species for rbcL, matK, and ITS2 combined**.

The evolutionary history was inferred using the Neighbor-Joining method (Saitou and Nei 1987). The optimal tree with the sum of branch length = 0.04111371 is shown. The tree is drawn to scale, with branch lengths in the same units as those of the evolutionary distances used to infer the phylogenetic tree. The evolutionary distances were computed using the Tamura 3-parameter method (Tamura 1992) and are in the units of the number of base substitutions per site. All positions containing gaps and missing data were eliminated. The rate variation among sites was modeled with a gamma distribution.

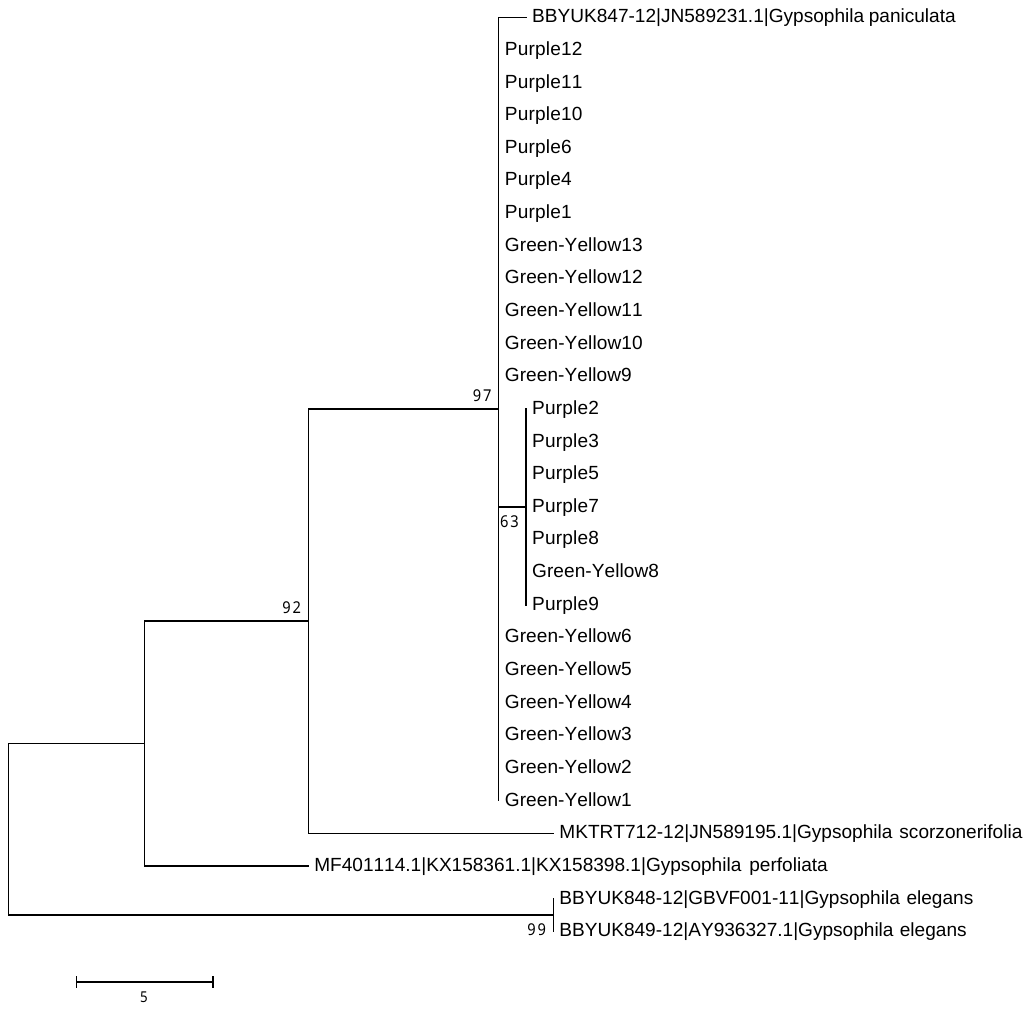

**Supplemental Figure 2.** **Maximum Parsimony analysis of baby’s breath color morphs in relation to other *Gypsophila* species for rbcL, matK, and ITS2 combined**.

The evolutionary history was inferred using the Maximum Parsimony method. Tree #1 out of 10 most parsimonious trees (length = 55) is shown. The consistency index is ( 0.930233), the retention index is ( 0.950820), and the composite index is 0.898957 (0.884483) for all sites and parsimony-informative sites (in parentheses). The percentage of replicate trees in which the associated taxa clustered together in the bootstrap test (500 replicates) are shown next to the branches (Felsenstein 1985). The MP tree was obtained using the Subtree-Pruning-Regrafting (SPR) algorithm with search level 1 in which the initial trees were obtained by the random addition of sequences (10 replicates). The tree is drawn to scale , with branch lengths calculated using the average pathway method and are in the units of the number of changes over the whole sequence. All positions containing gaps and missing data were eliminated.

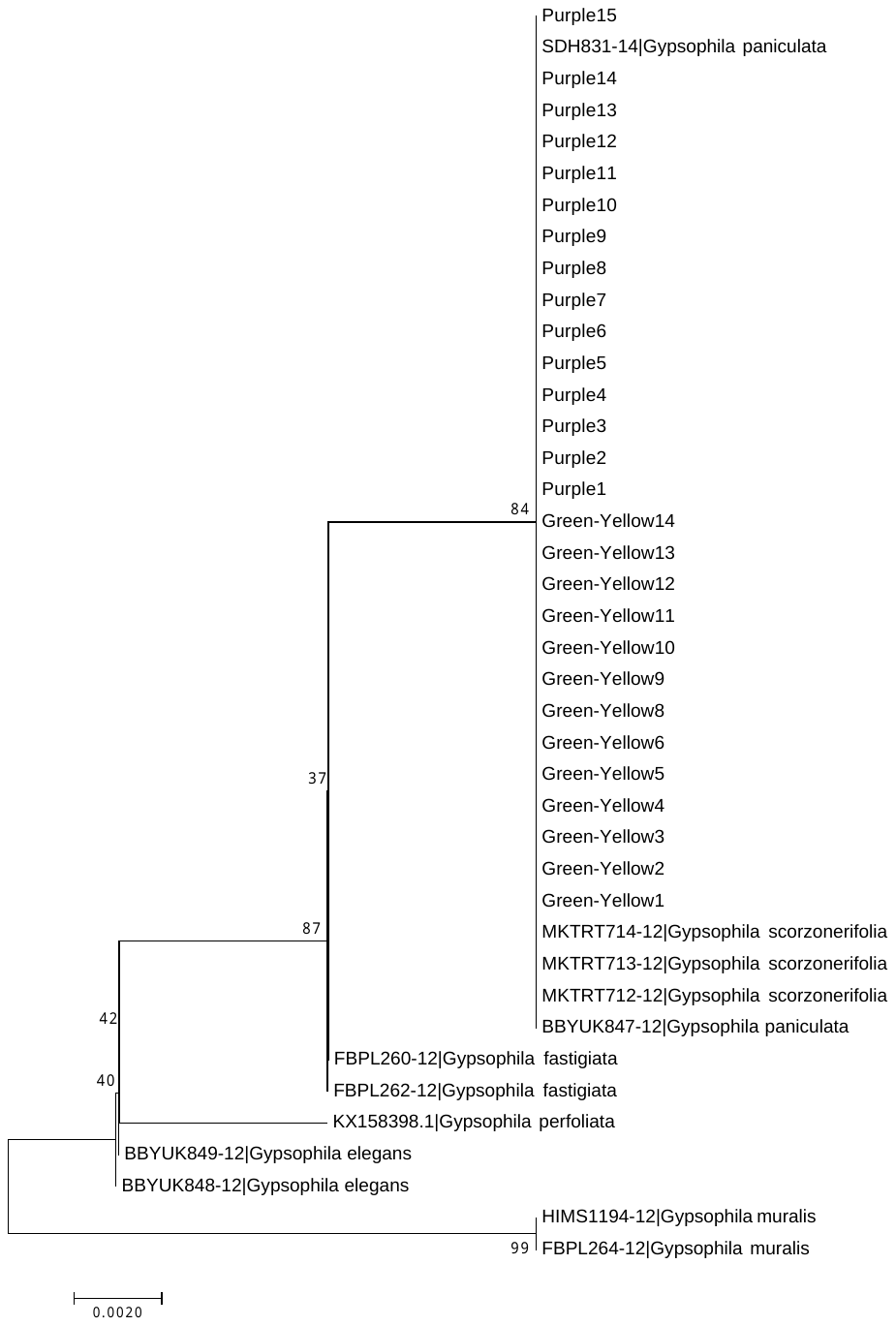

**Supplemental Figure 3:** **Neighbor joining analysis of baby’s breath color morphs in relation to other *Gypsophila* species for rbcL.**

The evolutionary history was inferred using the Neighbor-Joining method (Saitou and Nei 1987). The optimal tree with the sum of branch length = 0.0288485 is shown. The tree is drawn to scale, with branch lengths in the same units as those of the evolutionary distances used to infer the phylogenetic tree. The evolutionary distances were computed using the Jukes-Cantor method (Jukes and Cantor 1969) and are in the units of the number of base substitutions per site. All positions containing gaps and missing data were eliminated.

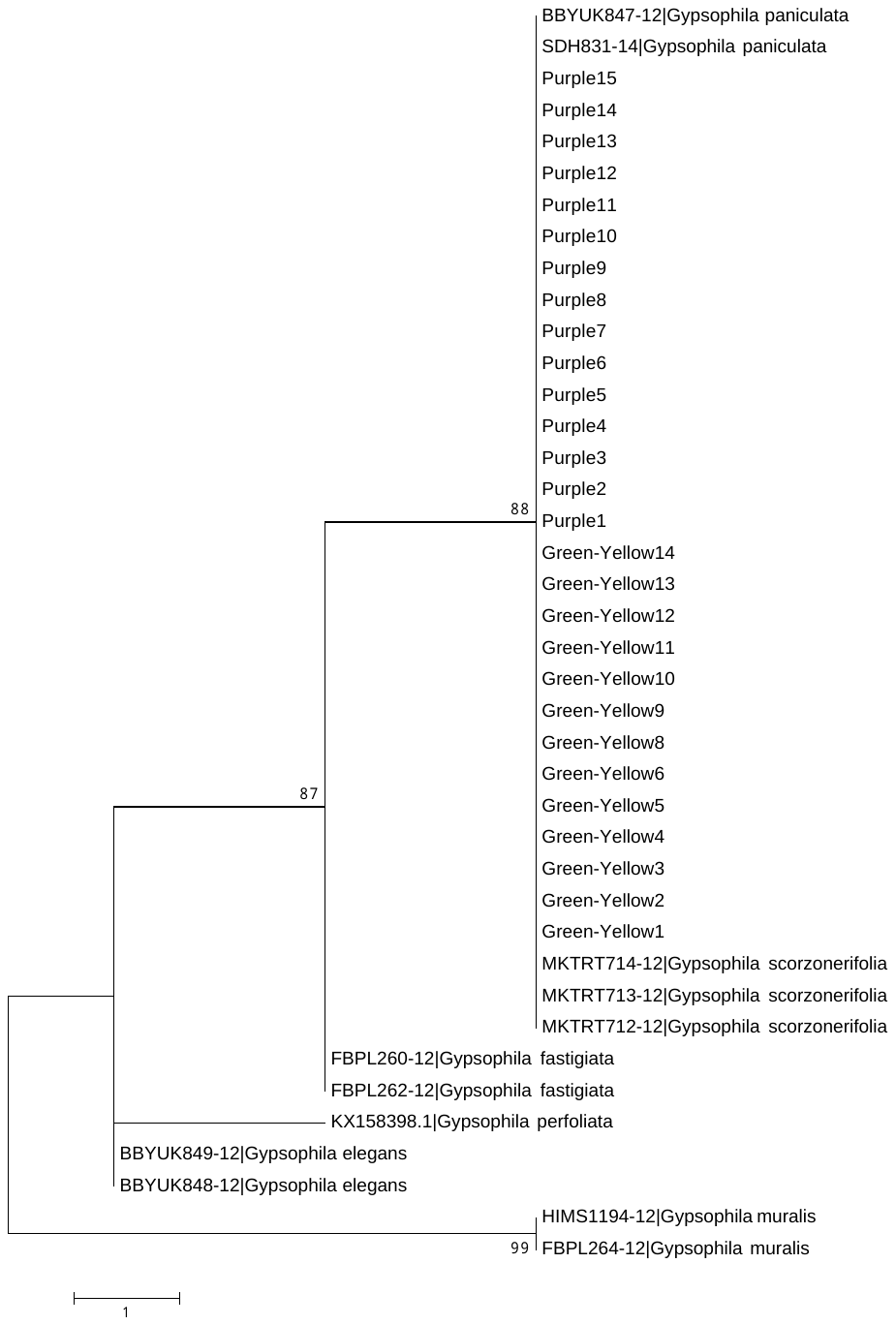

**Supplemental Figure 4:** **Maximum Parsimony analysis of baby’s breath color morphs in relation to other *Gypsophila* species for rbcL.**

The evolutionary history was inferred using the Maximum Parsimony method. Tree #1 out of 10 most parsimonious trees (length = 12) is shown. The consistency index is (1.000000), the retention index is (1.000000), and the composite index is 1.000000 (1.000000) for all sites and parsimony-informative sites (in parentheses). The percentage of replicate trees in which the associated taxa clustered together in the bootstrap test (500 replicates) are shown next to the branches (Felsenstein 1985). The MP tree was obtained using the Subtree-Pruning-Regrafting (SPR) algorithm (Nei and Kumar 2000) with search level 1 in which the initial trees were obtained by the random addition of sequences (10 replicates). The tree is drawn to scale, with branch lengths calculated using the average pathway method (Nei and Kumar 2000) and are in the units of the number of changes over the whole sequence. All positions containing gaps and missing data were eliminated.

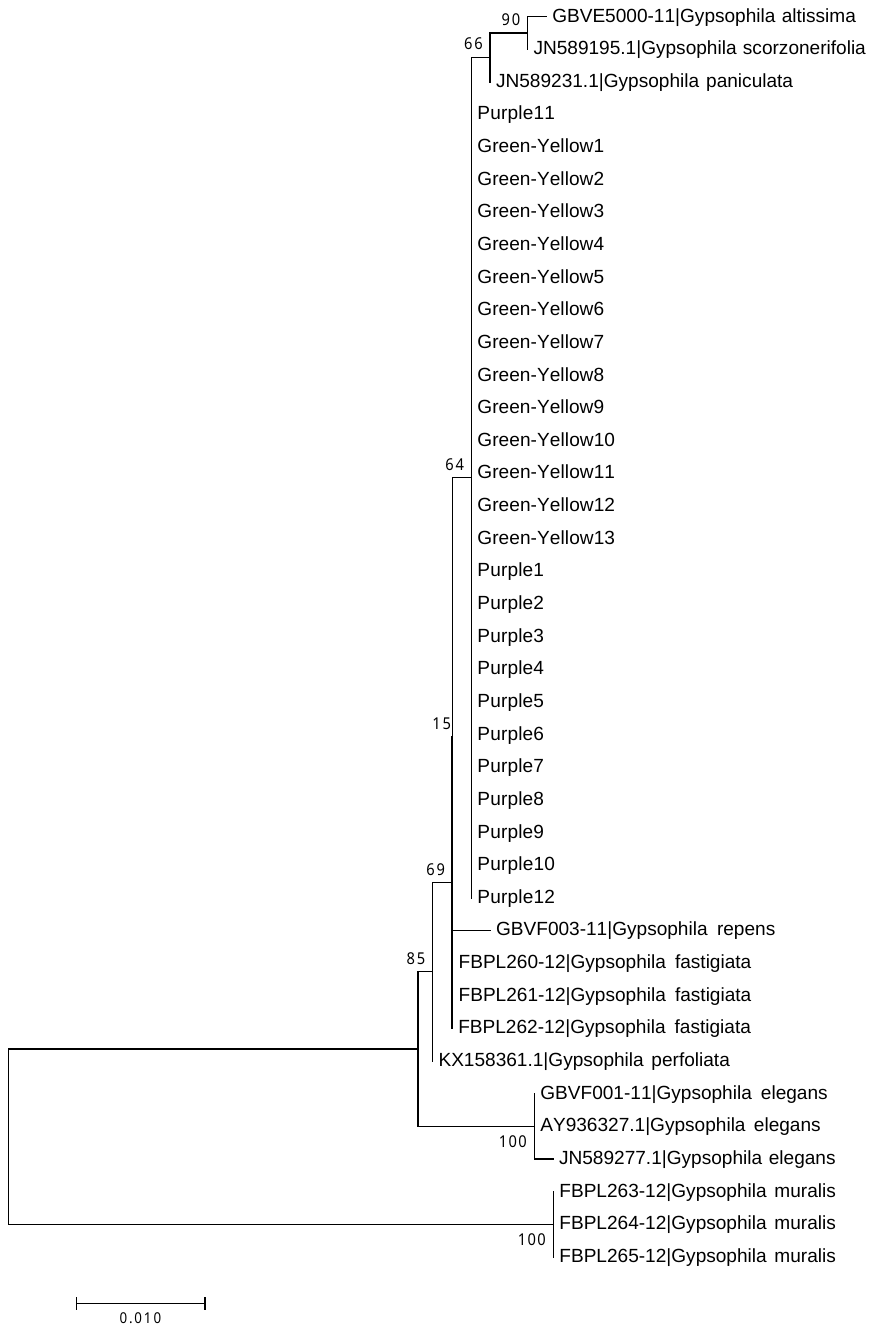

**Supplemental Figure 5:** **Neighbor joining analysis of baby’s breath color morphs in relation to other *Gypsophila* species for matK.**

The evolutionary history was inferred using the Neighbor-Joining method (Saitou an Nei 1987). The optimal tree with the sum of branch length = 0.0976890 is shown. The tree is drawn to scale, with branch lengths in the same units as those of the evolutionary distances used to infer the phylogenetic tree. The evolutionary distances were computed using the Tamura 3-parameter method (Tamura 1992) and are in the units of the number of base substitutions per site. All positions containing gaps and missing data were eliminated.

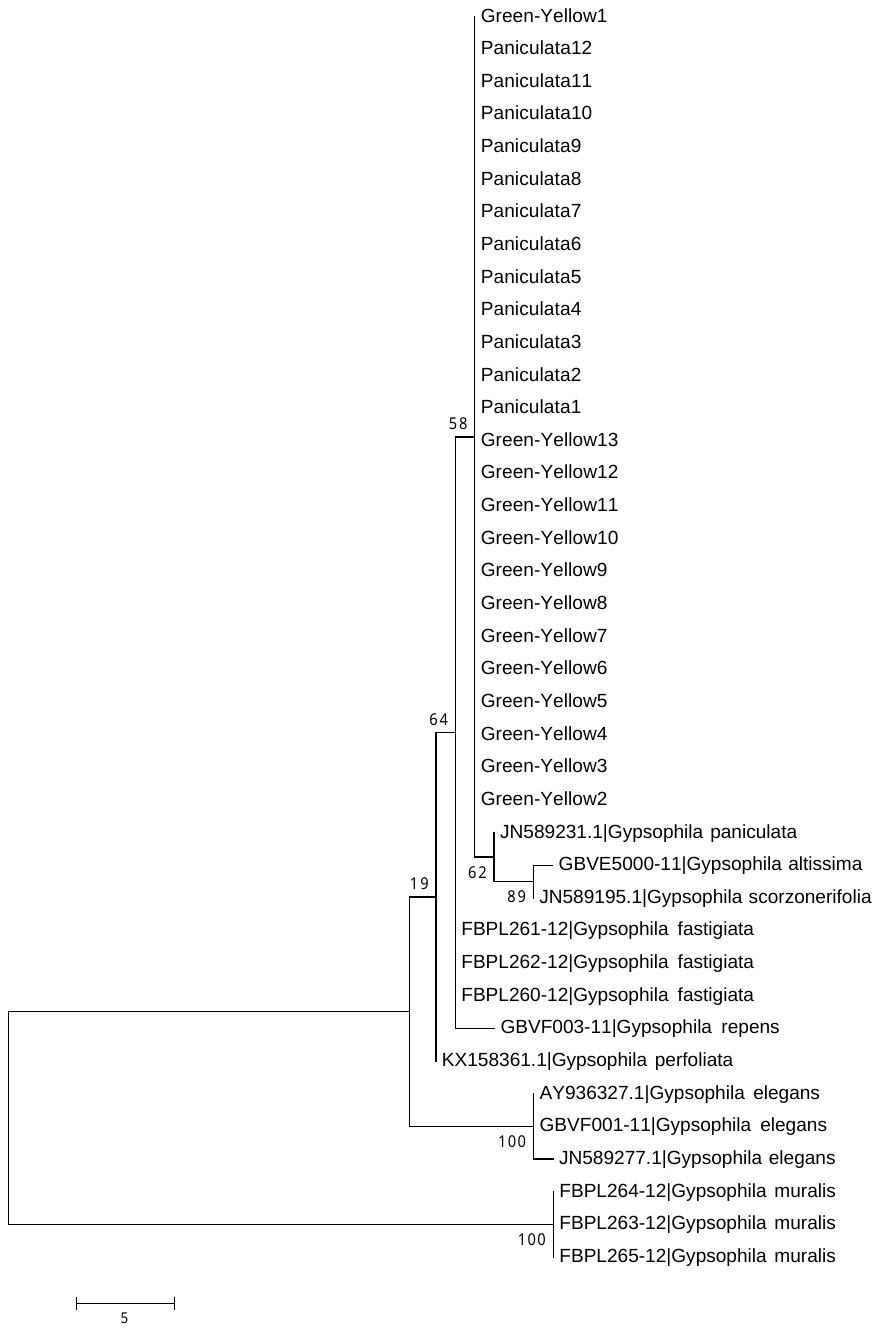

**Supplemental Figure 6:** **Maximum Parsimony analysis of baby’s breath color morphs in relation to other *Gypsophila* species for matK.**

The evolutionary history was inferred using the Maximum Parsimony method. Tree #1 out of 10 most parsimonious trees is shown (length = 64). The consistency index is (1.000000), the retention index is (1.000000), and the composite index is 1.000000 (1.000000) for all sites and parsimony-informative sites (in parentheses). The percentage of replicate trees in which the associated taxa clustered together in the bootstrap test (500 replicates) are shown next to the branches (Felsenstein 1985). The MP tree was obtained using the Subtree-Pruning-Regrafting (SPR) algorithm (Nei and Kumar 2000) with search level 1 in which the initial trees were obtained by the random addition of sequences (10 replicates). The tree is drawn to scale, with branch lengths calculated using the average pathway method (Nei and Kumar 2000) and are in the units of the number of changes over the whole sequence. All positions containing gaps and missing data were eliminated.

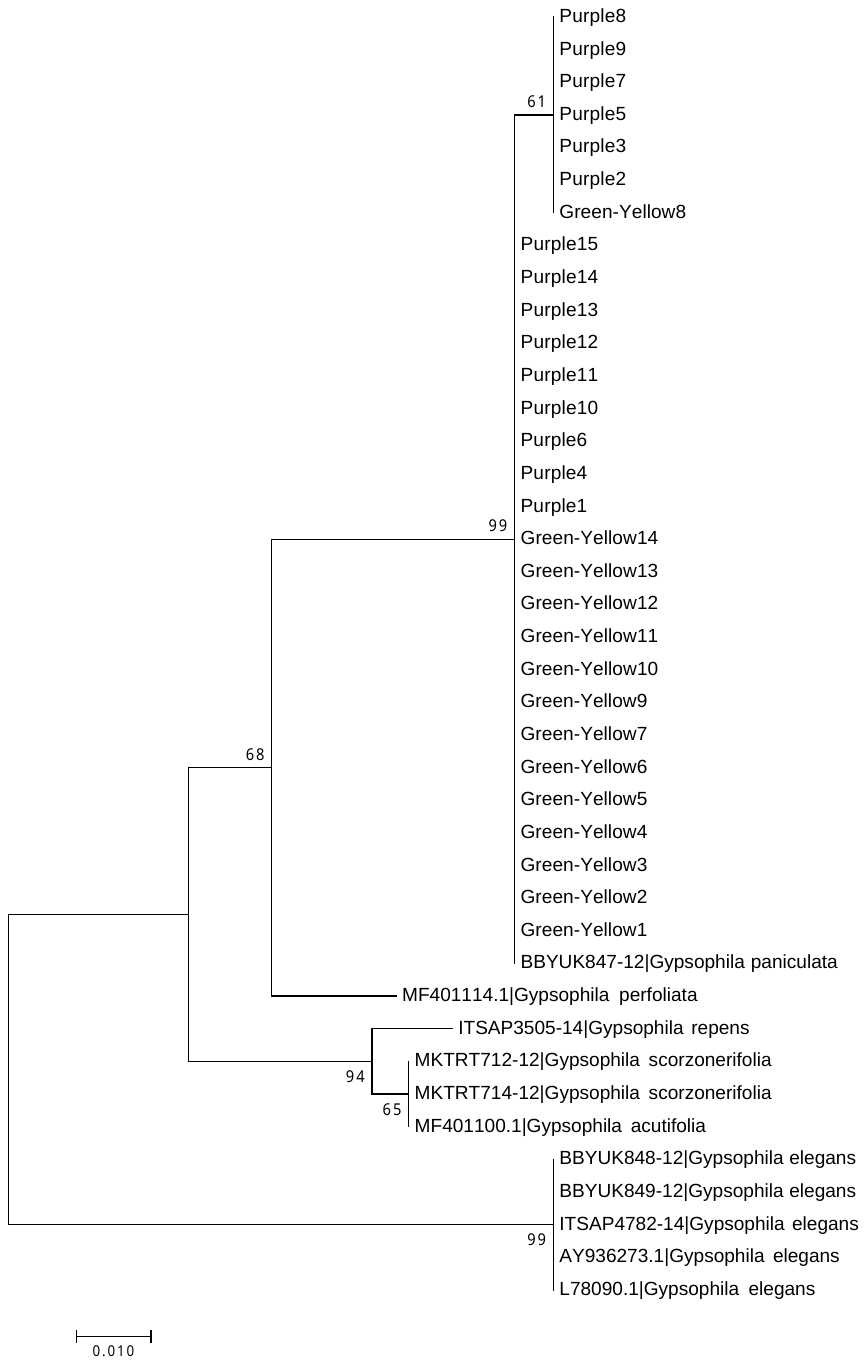

**Supplemental Figure 7:** **Neighbor joining analysis of baby’s breath color morphs in relation to other *Gypsophila* species for ITS2.**

The evolutionary history was inferred using the Neighbor-Joining method (Saitou and Nei 1987). The optimal tree with the sum of branch length = 0.20377523 is shown. The tree is drawn to scale, with branch lengths in the same units as those of the evolutionary distances used to infer the phylogenetic tree. The evolutionary distances were computed using the Jukes-Cantor method (Jukes and Cantor 1969) and are in the units of the number of base substitutions per site. All positions containing gaps and missing data were eliminated.

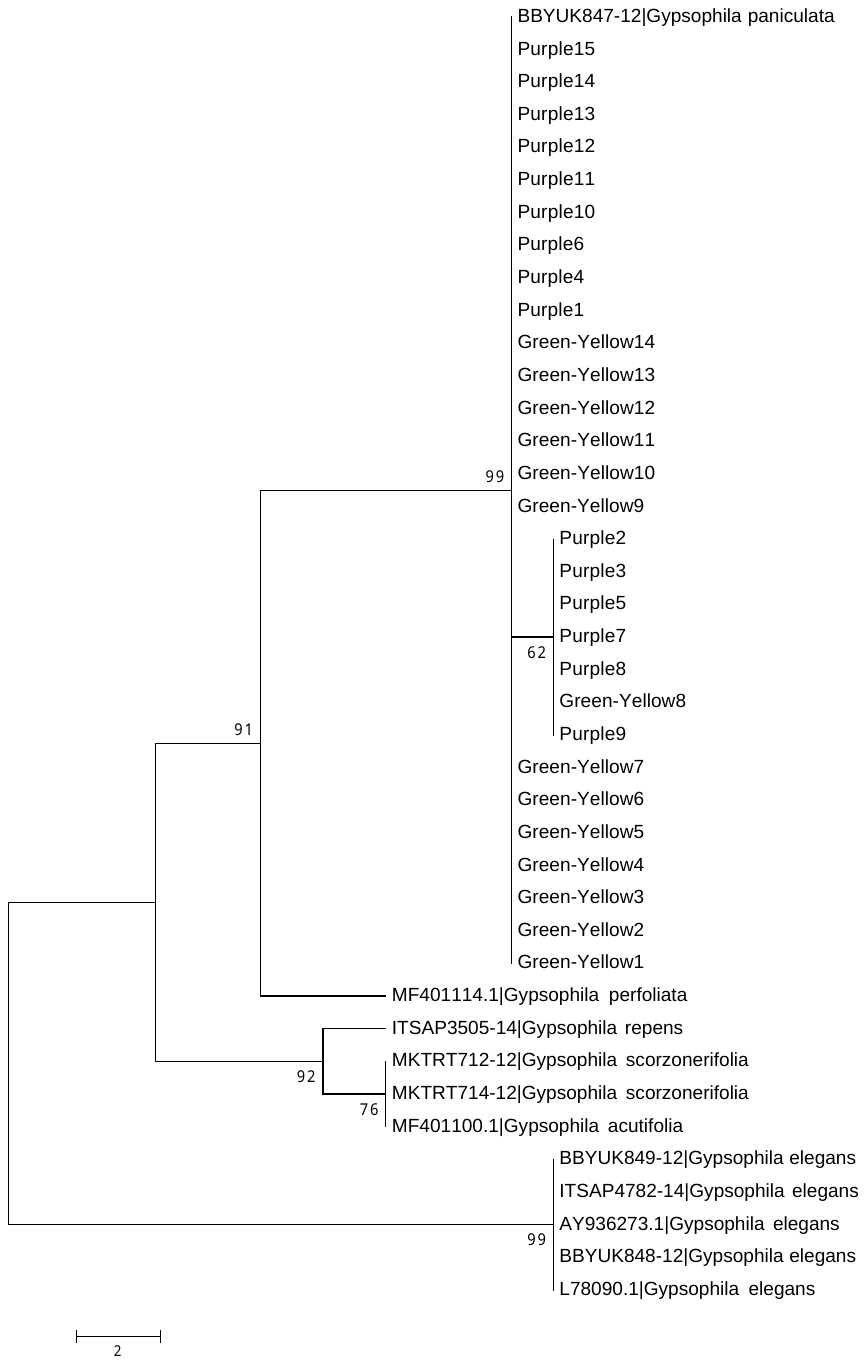

**Supplemental Figure 8:** **Maximum Parsimony analysis of baby’s breath color morphs in relation to other *Gypsophila* species for ITS2.**

The evolutionary history was inferred using the Maximum Parsimony method. Tree #1 out of 10 most parsimonious trees is shown (length = 36). The consistency index is (0.969697), the retention index is (0.993289), and the composite index is 0.965697 (0.963189) for all sites and parsimony-informative sites (in parentheses). The percentage of replicate trees in which the associated taxa clustered together in the bootstrap test (500 replicates) are shown next to the branches (Felsenstein 1985). The MP tree was obtained using the Subtree-Pruning-Regrafting (SPR) algorithm (Nei and Kumar 2000) with search level 1 in which the initial trees were obtained by the random addition of sequences (10 replicates). The tree is drawn to scale, with branch lengths calculated using the average pathway method (Nei and Kumar 2000) and are in the units of the number of changes over the whole sequence. All positions containing gaps and missing data were eliminated.
